# PerturbBridge: Conditional Latent Schrödinger Bridge for Single-Cell Perturbation Response Prediction

**DOI:** 10.64898/2026.09.24.754259

**Authors:** Zhihao Liu, Changzhi Jiang, Can Yang, Xiaoli Wang, Xiangrong Liu

**Author notes:** Corresponding author: Super Intelligent Medical Innovation Research Center, School of Informatics, Xiamen University, Xiamen 361005, Fujian, China.

## Abstract

**Motivation:** Single-cell perturbation prediction seeks to recover transcriptional responses to unseen interventions from unpaired control and perturbed populations. The Schrödinger Bridge provides a principled framework for modeling these transitions as stochastic transport, but learning bridge dynamics directly from high-dimensional, sparse gene-expression profiles remains challenging.

**Results:** We present PerturbBridge, a conditional latent Schrödinger Bridge framework for single-cell perturbation response prediction. A response-anchored autoencoder constructs a compact cellular representation that preserves dominant transcriptional response directions identified from training perturbations while encoding complementary cellular variation. In this latent space, PerturbBridge models conditional stochastic transport through a tractable approximation to the Schrödinger Bridge. We introduce the Perturbation-Induced Bridge Transformer, which uses perturbation- and time-dependent slots to guide interactions among latent coordinates during drift estimation. Evaluations on Norman, Sci-Plex3, and Replogle RPE1 demonstrate leading performance in differential-expression direction recovery and perturbation-effect correlation, alongside competitive gene-expression prediction accuracy.

**Availability:** The source code is available at https://github.com/echQoQ/PerturbBridge, with an archived version at https://doi.org/10.5281/zenodo.24272128.

## 1. Introduction

Single-cell perturbation technologies, including CRISPR-based screens and chemical treatment assays, enable systematic investigation of transcriptional responses to genetic and pharmacological interventions (Dixit et al., 2016; Srivatsan et al., 2020). Beyond advancing our understanding of gene regulation and drug action, these technologies provide valuable data for developing AI-driven virtual cell models that seek to predict cellular behavior under different conditions (Bunne et al., 2024). A central challenge toward this goal is predicting responses to unseen perturbations, as the vast space of possible interventions, dosages, and cellular contexts makes exhaustive experimental characterization impractical (Wei et al., 2026). This task is further complicated by the destructive nature of single-cell sequencing, which prevents direct observation of the same cell before and after perturbation. Consequently, control and perturbed cells are observed as unpaired populations (Bunne et al., 2023), requiring computational models to recover both transcriptional response patterns and population-level variability from these observations.

A variety of computational methods have been developed for single-cell perturbation prediction. GEARS (Roohani et al., 2024) exploits gene relationships to predict transcriptional responses to genetic perturbations, while scGPT (Cui et al., 2024) leverages pretrained single-cell representations for perturbation modeling. Conditional generative models, such as chemCPA (Hetzel et al., 2022) and PRnet (Qi et al., 2024), predict responses to chemical perturbations, whereas PerturbNet (Yu et al., 2025) employs conditional normalizing flows to model perturbed cell distributions. Recent approaches have further explored generative dynamics for perturbation prediction. CellFlow (Klein et al., 2025) learns perturbation-conditioned transport through flow matching, while scLDM (Wu et al., 2026) uses conditional latent diffusion to generate post-perturbation cellular states. These advances have extended perturbation prediction toward modeling the distributional changes induced by interventions.

The Schrödinger Bridge (SB) provides a principled framework for stochastic transport between probability distributions, seeking a stochastic process that satisfies prescribed endpoint marginals while minimizing relative entropy with respect to a reference process (Léonard, 2013). For general endpoint distributions, however, the SB problem typically lacks a closed-form solution, and numerical methods often involve computationally demanding iterative procedures. Recent advances have developed practical approximations for learning SB dynamics (De Bortoli et al., 2021; Shi et al., 2023; Liu et al., 2023). As SB formulates transport at the distribution level, it does not require observed cell-level correspondences, making it particularly suitable for single-cell perturbation prediction, where control and perturbed cells are observed as unpaired populations. However, applying SB directly to high-dimensional, sparse gene-expression data remains challenging. Departures (Chi et al., 2026) approaches this problem by constructing approximate endpoint couplings through minibatch optimal transport and jointly learning two coupled bridges for continuous gene expression and discrete gene activation states. This design explicitly accounts for expression sparsity, while requiring the learning of two stochastic processes at the gene level. An alternative is to learn bridge dynamics in a compact cellular latent space, potentially capturing coordinated transcriptional variation in fewer dimensions. The challenge then lies in preserving the perturbation-induced transcriptional changes needed for accurate population-level prediction.

To address these challenges, we propose PerturbBridge, a conditional latent Schrödinger Bridge framework for predicting transcriptional responses to unseen perturbations from unpaired single-cell observations. PerturbBridge first learns a compact cellular representation using a response-anchored autoencoder (RA-AE), which explicitly preserves dominant transcriptional response directions identified from training perturbations while encoding the remaining cellular variation. Within this representation, PerturbBridge learns a conditional latent SB using a tractable bridge-matching formulation. To model how cellular states and perturbation conditions influence the bridge dynamics, we develop the Perturbation-Induced Bridge Transformer (PIBT), which uses perturbation- and time-dependent representations to guide interactions among latent features during conditional drift estimation. During inference, the learned stochastic dynamics transport encoded control cells toward perturbation-specific latent populations, which are subsequently decoded into gene-expression profiles.

We evaluate PerturbBridge on three benchmarks spanning genetic and chemical perturbations: Norman (Norman et al., 2019), Sci-Plex3 (Srivatsan et al., 2020), and Replogle RPE1 (Replogle et al., 2022). Across these datasets, PerturbBridge demonstrates strong performance in gene-expression prediction, differential-expression recovery, and perturbation discrimination. In particular, it achieves leading results in differential-expression direction recovery and perturbation-effect correlation across all three benchmarks, while maintaining high gene-expression prediction accuracy. Ablation studies further support the contributions of response-anchored representation learning and PIBT-based conditional drift modeling, including the benefits of condition-guided interactions among latent coordinates.

## 2. Methods and materials

### 2.1. Overview of PerturbBridge

Let *x* ∈ ℝ ^*G*^ denote the gene-expression profile of a single cell, *τ* its cellular context, and *c* a perturbation condition comprising the perturbation identity and associated treatment covariates. We denote the control and perturbed cell distributions by 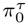 and 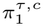, respectively. Since single-cell RNA sequencing is destructive, these populations are observed without cell-level correspondence. Given a control population and a specified perturbation condition, our objective is to predict the corresponding perturbed cell distribution, including for perturbations unseen during training.

PerturbBridge models the transition between control and perturbed populations as a conditional Schrödinger Bridge (SB) in a compact cellular latent space, as illustrated in Fig. 1. The latent representation is constructed to preserve dominant transcriptional response directions identified from training perturbations while encoding the remaining expression variation. We achieve this through a response-anchored autoencoder (RA-AE), which provides a shared latent space for control and perturbed cells. Stochastic transport between the resulting latent distributions is then learned through a tractable bridge-matching formulation. To account for the dependence of bridge dynamics on cellular states, time, and perturbation conditions, we introduce a condition-guided interaction mechanism that coordinates information exchange among latent features. This mechanism is implemented in the Perturbation-Induced Bridge Transformer (PIBT) to parameterize the conditional drift field.

**Figurxe 1.**
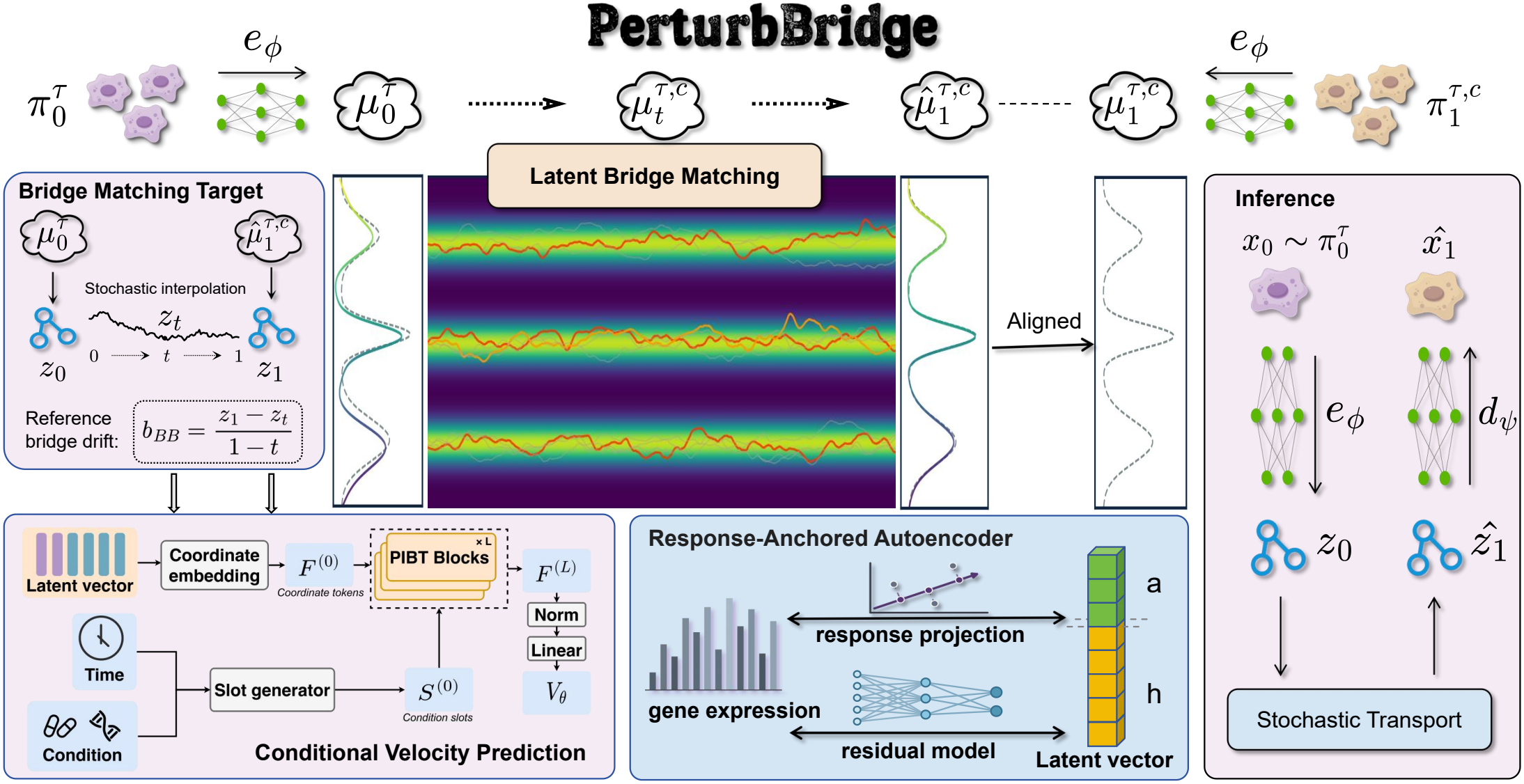
Overview of PerturbBridge. PerturbBridge models unpaired single-cell perturbation responses through conditional stochastic transport in a compact latent space. A response-anchored autoencoder (RA-AE) first learns cellular representations that preserve dominant perturbation-response directions. In the learned latent space, stochastic interpolation between control and perturbed samples provides reference bridge trajectories for training. A perturbation-induced bridge transformer (PIBT) predicts the conditional drift field from the intermediate latent state, bridge time, and perturbation condition. During inference, control cells are encoded, transported through the learned stochastic dynamics, and decoded to generate the predicted perturbed cell population.

Training proceeds in two stages. RA-AE is first optimized for gene-expression reconstruction, after which its encoder and decoder are frozen. The conditional bridge model is then trained using latent representations of unpaired control and perturbed cells. During inference, control cells are encoded into the latent space and transported under the learned stochastic dynamics conditioned on the target perturbation. The resulting latent states are decoded to generate the predicted perturbed cell population.

### 2.2 Cellular Representation

To support stochastic transport in the latent space, we construct a cellular representation that explicitly retains major perturbation-associated transcriptional response directions. We implement this design through a response-anchored autoencoder (RA-AE), which combines fixed response-aligned coordinates with a nonlinear representation of residual expression variation.

We construct a shared response subspace from perturbation-induced expression changes. For each cellular context *τ* and perturbation condition *c*, the transcriptional response is estimated as the difference between the corresponding perturbed and control population means:

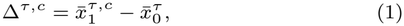

where 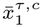 and 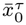 denote the mean expression profiles of perturbed and control cells, respectively. Stacking these response vectors as columns yields Δ ∈ ℝ ^*G×M*^, where *M* is the number of context– condition pairs. We perform singular value decomposition Δ = *L*Σ*V* ^*⊤*^ and retain the leading *r* left singular vectors to form an orthonormal basis *U* ∈ ℝ ^*G×r*^, satisfying *U*^*⊤*^*U* = *I*_*r*_. This basis spans the dominant transcriptional response directions across perturbation conditions.a

Given a gene-expression profile *x*, we center it using the overall expression mean *µ* ∈ ℝ ^*G*^ and decompose the centered profile into response-aligned and orthogonal components:

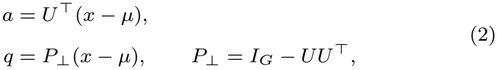

where *a* ∈ ℝ ^*r*^ contains the response-aligned coordinates and *q* ∈ ℝ ^*G*^ is the orthogonal residual. RA-AE retains *a* directly and maps the residual to a lower-dimensional representation *h* = *E*_*ϕ*_(*q*) ∈ ℝ ^*d−r*^ through a nonlinear encoder. The resulting latent representation is 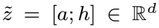. Reconstruction combines the response-aligned component with the decoded residual:

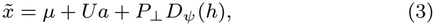

where *D*_*ψ*_ denotes the residual decoder. The residual encoder and decoder are trained by minimizing the mean squared reconstruction error:

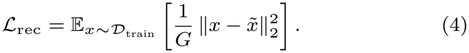

After representation learning, the latent representations are normalized for subsequent bridge training. Let *m* ∈ ℝ ^*d*^ denote the mean latent vector and *s >* 0 a global scale defined by

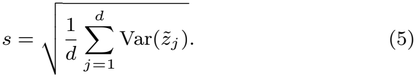

The complete encoder and decoder are then defined as

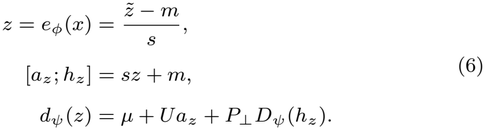

By construction, *U*^*⊤*^*P*_*⊥*_ = 0 and *U*^*⊤*^*U* = *I*_*r*_, yielding

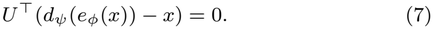

Thus, RA-AE preserves the response-aligned coordinates exactly during reconstruction, with reconstruction errors confined to the orthogonal complement.

### 2.3. Conditional Latent Schrödinger Bridge

Using the learned cellular representation, we formulate perturbation prediction as conditional stochastic transport in the latent space. The fixed encoder *e*_*ϕ*_ induces the control and perturbed latent distributions

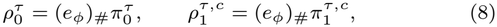

where (*e*_*ϕ*_)_#_ denotes the pushforward operator. Our objective is to learn a stochastic process connecting 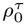 to 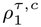.

The Schrödinger Bridge (SB) formulates stochastic transport as relative entropy minimization over path measures subject to prescribed endpoint distributions (Léonard, 2013). Given a reference path measure ℝ ^*τ*^, the conditional SB problem is

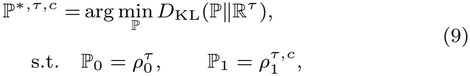

where ℙ _*t*_ denotes the time-*t* marginal distribution. By the relative entropy decomposition, the SB solution can be represented as a mixture of conditional reference paths:

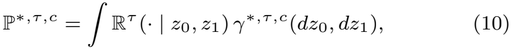

where the optimal endpoint coupling satisfies

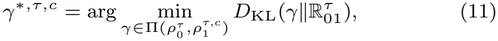

with 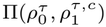 denoting the set of admissible couplings and 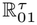 the endpoint joint distribution of the reference process.

To obtain a tractable formulation, we choose a Brownian reference process *dZ*_*t*_ = *σ*_B_ *dW*_*t*_ with constant, isotropic diffusion coefficient *σ*_B_ *>* 0, initialized from 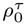. Instead of solving for the optimal coupling, we independently sample endpoints from the control and perturbed populations, corresponding to 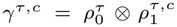. This avoids explicit coupling optimization while preserving the prescribed endpoint marginals, although the resulting transport need not be entropy-optimal.

Following the analytic conditional bridge construction of I^2^SB (Liu et al., 2023), the intermediate state at *t* ∈ (0, 1), conditioned on an endpoint pair (*z*_0_, *z*_1_) ~ *γ*^*τ,c*^, follows

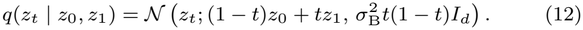

This distribution permits direct sampling:

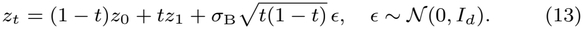

The corresponding endpoint-conditioned reference drift is

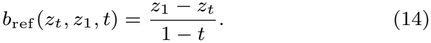

We learn a conditional drift field *v*_*θ*_(*z*_*t*_, *t, ĉ*), where *ĉ* = *κ*_*ω*_(*τ, c*) represents the perturbation identity, associated covariates, and cellular context. Following bridge matching (Shi et al., 2023), we minimize

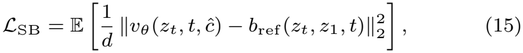

where the expectation is over cellular contexts, perturbation conditions, endpoint pairs, bridge times, and Gaussian noise.

The population minimizer of this objective is the conditional expectation of the reference drift:

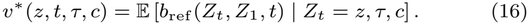

By Markovian projection, a diffusion with drift *v*^*\**^ and diffusion coefficient *σ*_B_ shares the time marginals of the conditional bridge mixture. Specifically, the marginal density 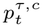 satisfies

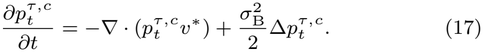

Under suitable regularity and uniqueness conditions, this diffusion, initialized from 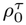, recovers the prescribed endpoint distributions in the continuous-time limit. PerturbBridge parameterizes the conditional drift using the Perturbation-Induced Bridge Transformer (PIBT), described next.

### 2.4. Perturbation-Induced Bridge Transformer

Learning the conditional bridge requires a drift field that responds to the evolving cellular state, bridge time, and perturbation condition. Since perturbation responses may involve coordinated changes across latent dimensions, the drift predictor should capture condition-dependent interactions among these coordinates. Rather than relying solely on feature modulation for conditioning (Peebles and Xie, 2023), we develop the Perturbation-Induced Bridge Transformer (PIBT), which uses a compact set of perturbation- and time-conditioned slots to mediate information exchange among latent coordinates. The architecture is illustrated in Fig. 2.

**Figure 2.**
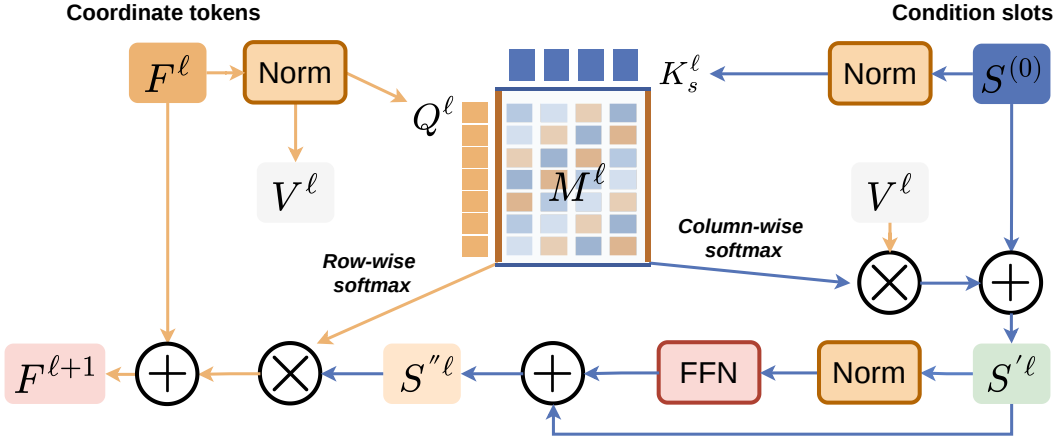
Architecture of the Perturbation-Induced Bridge Transformer (PIBT) block.

Given an intermediate latent state *z*_*t*_ ∈ ℝ ^*d*^, PIBT represents each latent coordinate as an individual token through feature-wise tokenization:

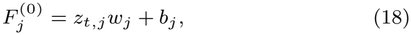

where *w*_*j*_, *b*_*j*_ ∈ ℝ ^*H*^ are coordinate-specific learnable parameters, yielding *F* ^(0)^ ∈ ℝ ^*d×H*^. To incorporate perturbation and temporal information, PIBT generates *K* condition-induced slots. Let *e*_*t*_ denote the learned time embedding and *e*_*c*_ = *W*_*c*_*ĉ*+ *b*_*c*_ the projected condition embedding. The slots are initialized as

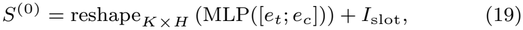

where *I*_slot_ ∈ ℝ ^*K×H*^ contains learnable slot identity embeddings. These slots provide condition-dependent representations through which latent coordinates can exchange information.

To mediate interactions between coordinate tokens and condition-induced slots, we build on the shared-affinity mechanism of BiXT (Hiller et al., 2024) and organize information exchange into a sequential read–process–write procedure. At block *l*, the coordinate– slot affinity matrix is computed as

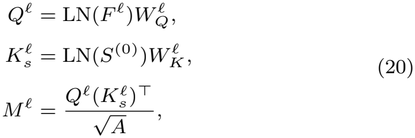

where *M*^*l*^ ∈ ℝ ^*d×K*^ and *A* is the affinity projection dimension. The affinity scores jointly depend on the current coordinate representations and the condition-induced slots, enabling state-dependent interactions under the specified perturbation and bridge time.

A shared affinity matrix determines attention weights in both directions:

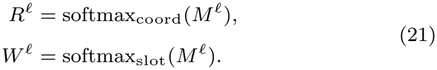

Here, softmax_coord_ normalizes across coordinate tokens for each slot, determining how information is aggregated into the slots. Conversely, softmax_slot_ normalizes across slots for each coordinate, determining how information is distributed back to the coordinate tokens. Both directions therefore share the same underlying coordinate–slot affinity scores.

During the read stage, coordinate information is aggregated into the slots and processed through a slot-wise feed-forward network:

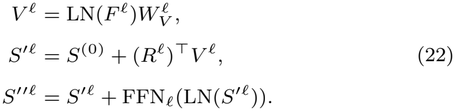

The processed slot representations are then written back to the coordinate tokens:

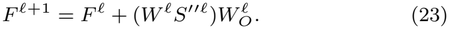

This read–process–write procedure enables information exchange among latent coordinates through the compact slot representations, without direct coordinate-wise self-attention. Across blocks, the same condition-induced slots *S*^(0)^ are reused, while the coordinate representations are progressively updated. Consequently, the coordinate–slot affinities are recomputed at each block, allowing the interactions to adapt to the evolving cellular state.

After *L* blocks, a shared prediction head maps each coordinate representation to its corresponding drift component:

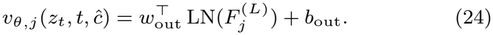

The resulting drift vector *v*_*θ*_(*z*_*t*_, *t, ĉ*) ∈ ℝ ^*d*^ parameterizes the conditional stochastic dynamics introduced in Section 2.3.

### 2.5. Stochastic Inference

Given a control cell 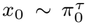 and a target perturbation condition *c*, PerturbBridge generates the corresponding perturbed cell state through the learned conditional stochastic dynamics. The initial latent state and condition embedding are obtained as *z*_0_ = *e*_*ϕ*_(*x*_0_) and *ĉ* = *κ*_*ω*_(*τ, c*), respectively. The latent evolution follows the stochastic differential equation

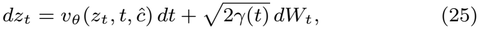

where *v*_*θ*_ denotes the conditional drift predicted by PIBT and *W*_*t*_ is standard Brownian motion. To maintain consistency with the reference diffusion used for bridge matching, we set 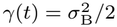.

We numerically integrate the stochastic dynamics using the Euler– Maruyama scheme (Higham, 2001):

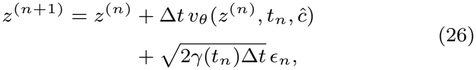

where *z*^(0)^ = *z*_0_, *t*_*n*_ = *n/N*, Δ*t* = 1*/N*, and *ϵ*_*n*_ ~ *N*(0, *I*_*d*_) are independently sampled for *n* = 0, …, *N* − 1. After *N* integration steps, the terminal latent state is mapped back to gene-expression space using the pretrained RA-AE decoder:

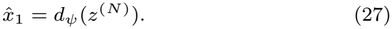

Repeating this procedure across control cells and independently sampled stochastic trajectories produces the predicted perturbed cell population under condition *c*.

### 2.6. Datasets

We evaluate PerturbBridge on three single-cell perturbation benchmarks spanning genetic and chemical interventions: Norman (Norman et al., 2019), Sci-Plex3 (Srivatsan et al., 2020), and Replogle RPE1 (Replogle et al., 2022). We use the datasets released with PerturbNet (Yu et al., 2025) for Norman and Sci-Plex3, and the preprocessed Replogle RPE1 dataset distributed through GEARS (Roohani et al., 2024). Norman comprises 109,738 cells and 2,279 genes, covering single-gene and combinatorial CRISPR perturbations, with 95 of 236 perturbation conditions held out for testing. Sci-Plex3 contains 648,737 cells and 5,087 genes, spanning 188 drugs across three cell lines and four dosage levels, with 38 drugs reserved for testing. Replogle RPE1 comprises 162,733 cells and 5,000 genes, covering 1,543 genetic perturbation targets, of which 145 are held out for testing. All splits are defined at the perturbation level, with separate validation subsets drawn from the training perturbations for representation and bridge learning.

### 2.7. Implementation Details

PerturbBridge is implemented in PyTorch and trained in two stages. RA-AE employs an encoder with hidden dimensions of 512 and 256 and a symmetric decoder, using GELU activations. The latent and response dimensions (*d, r*) are set to (24, 4) for Norman and Sci-Plex3 and (64, 8) for Replogle RPE1. For PIBT, the hidden dimension, number of slots, number of blocks, and affinity dimension (*H, K, L, A*) are set to (128, 4, 3, 32) for Norman and (256, 8, 4, 64) for Sci-Plex3 and Replogle RPE1. RA-AE is trained using Adam (Kingma and Ba, 2014) with a batch size of 4,096 and an initial learning rate of 5 × 10^*−*4^, after which its parameters are frozen. PIBT is optimized using AdamW (Loshchilov and Hutter, 2017) with a weight decay of 10^*−*4^. The batch sizes are 2,048 for Norman and Replogle RPE1 and 4,096 for Sci-Plex3, with initial learning rates of 10^*−*4^ and 5 × 10^*−*5^, respectively. Both stages are trained for 200 epochs using cosine learning rate schedules. For chemical perturbations, we use precomputed 384-dimensional embeddings from ChemBERTa (Chithrananda et al., 2020), derived from molecular SMILES representations, together with dose and cell-line covariates. For genetic perturbations, gene embeddings are constructed by applying PCA to Gene Ontology (GO) annotations (Aleksander et al., 2023). The resulting embeddings have dimensions of 256 for Norman and 384 for Replogle RPE1, with individual gene embeddings summed for combinatorial perturbations. The pretrained chemical representations and GO-derived embeddings remain fixed, while the downstream condition encoders are optimized jointly with PIBT. The reference diffusion scale is set to *σ*_B_ = 1.0, and stochastic inference is performed using 100 Euler–Maruyama integration steps. All experiments are conducted in FP32 on a single NVIDIA GeForce RTX 3090 GPU.

## 3. Experiment

### 3.1. Evaluation Protocol

We compare PerturbBridge with representative single-cell perturbation prediction methods. For the genetic perturbation benchmarks, Norman and Replogle RPE1, the baselines include Control, CPA (Lotfollahi et al., 2023), GEARS (Roohani et al., 2024), PerturbNet (Yu et al., 2025), scGPT (Cui et al., 2024), CellFlow (Klein et al., 2025), Departures (Chi et al., 2026), and scLDM (Wu et al., 2026). For Sci-Plex3, we compare against Control, PRnet (Qi et al., 2024), chemCPA (Hetzel et al., 2022), PerturbNet, CellFlow, Departures, and scLDM.

We evaluate expression accuracy, differential-expression recovery, and perturbation discrimination using selected metrics from Cell-Eval (Adduri et al., 2026). Expression accuracy is measured using mean squared error (MSE) and mean absolute error (MAE) on pseudobulk expression profiles. Differential-expression recovery is assessed using DE Direction Match (DirM) and Spearman correlation of log-fold changes (SpLFC), which evaluate agreement in differential-expression directions and gene-level response rankings, respectively. Pearson Δ correlation measures the agreement of perturbation-induced expression changes relative to controls, while Perturbation Discrimination Scores (PDS) evaluate perturbation discrimination using *L*_1_, *L*_2_, and cosine distances.

### 3.2. Main Results: Performance on Unseen Perturbations

#### Overall performance across datasets

We evaluate PerturbBridge on three established single-cell perturbation benchmarks—Norman, Sci-Plex3, and Replogle RPE1—spanning both genetic and chemical perturbations. We compare PerturbBridge with representative baselines encompassing direct expression prediction and conditional generative modeling, using metrics that assess expression accuracy, differential-expression recovery, and perturbation discrimination. The overall results are presented in Fig. 3(a–c).

**Figure 3.**
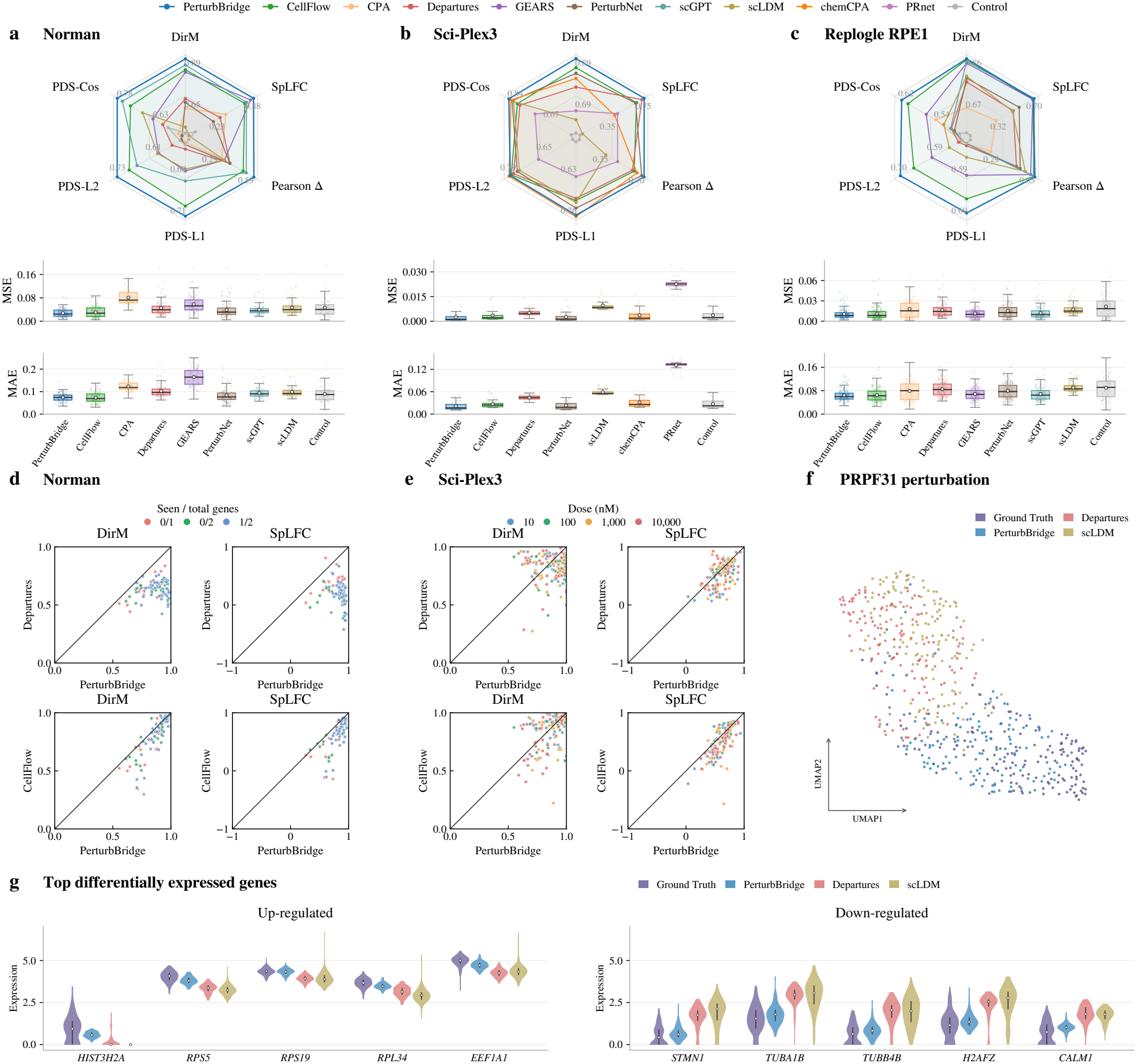
Evaluation of PerturbBridge across genetic and chemical perturbation benchmarks. (a–c) Performance comparison on Norman, Sci-Plex3, and Replogle RPE1. Radar plots summarize six perturbation-response metrics, while box plots show the distributions of MSE and MAE across test conditions. (d) Perturbation-level comparisons on Norman, stratified by the proportion of previously seen target genes. (e) Dose-stratified comparisons on Sci-Plex3. In (d) and (e), each point represents a test condition, comparing PerturbBridge against Departures or CellFlow using DirM and SpLFC. (f) UMAP visualization of ground-truth and predicted cells under PRPF31 perturbation in Replogle RPE1. (g) Expression distributions of the top five upregulated and downregulated genes under PRPF31 perturbation, comparing PerturbBridge with Departures, scLDM, and ground truth.

Among the generative baselines, CellFlow, a flow matching-based approach, exhibits strong and relatively consistent performance across the three datasets, approaching PerturbBridge on Replogle RPE1. However, larger performance gaps emerge on Norman and Sci-Plex3, particularly in differential-expression recovery. PerturbNet, based on a conditional invertible neural network (cINN), performs competitively on Sci-Plex3 but shows weaker performance on Replogle RPE1, whereas the latent diffusion-based scLDM exhibits limited recovery of perturbation-induced transcriptional changes, particularly on Norman and Sci-Plex3. Departures provides a particularly relevant comparison, as it also builds on the Schrödinger Bridge framework but employs dual bridges in gene-expression space with minibatch optimal transport for endpoint coupling. Despite this shared theoretical foundation, PerturbBridge achieves consistently stronger perturbation-response prediction, with particularly pronounced advantages in differential-expression recovery on Norman and perturbation discrimination on Replogle RPE1. These findings further support the effectiveness of latent-space bridge modeling.

Overall, PerturbBridge achieves state-of-the-art performance across the three benchmarks, with particularly strong results in recovering perturbation-induced transcriptional changes. It consistently achieves the highest DirM and Pearson Δ scores across all three datasets, indicating accurate recovery of differential-expression directions and expression changes relative to controls. Its leading SpLFC scores on Norman and Sci-Plex3 further demonstrate strong agreement in the relative ordering of gene-level log-fold changes. PerturbBridge also achieves the highest PDS scores across all three distance variants on Norman and Replogle RPE1, reflecting improved discrimination of perturbation-specific transcriptional signatures. Notably, these advantages are accompanied by the lowest MSE across all three benchmarks, indicating that improved perturbation-response fidelity is achieved without compromising overall expression prediction accuracy.

#### Consistency across perturbation conditions

To assess the consistency of performance improvements across individual perturbation conditions, we compare PerturbBridge with CellFlow and Departures using condition-level DirM and SpLFC scores (Fig. 3d–e). DirM measures agreement in the direction of differential expression between predicted and observed responses, whereas 0.5213 0.7200 0.0298 0.0757 SpLFC evaluates the Spearman correlation of gene-level log-fold changes, reflecting the recovery of relative transcriptional response patterns. On Norman, PerturbBridge outperforms both baselines across nearly all perturbation conditions on both metrics, with Departures exhibiting the largest performance gaps. These advantages are observed across different levels of target-gene novelty, including single-gene and combinatorial perturbations involving unseen targets. On Sci-Plex3, although the performance differences are generally smaller, PerturbBridge achieves higher scores for the majority of drug–dose conditions, with a particularly consistent advantage in SpLFC. Together, these results indicate that the improvements in differential-expression recovery extend across diverse perturbation conditions rather than being driven by a limited number of favorable cases.

#### Transcriptional response fidelity

To further examine the fidelity of predicted transcriptional responses, we investigate PRPF31 perturbation in Replogle RPE1 (Fig. 3f–g). In the UMAP visualization, PerturbBridge-predicted cells show closer alignment with the observed perturbed population, whereas predictions from Departures and scLDM exhibit more pronounced spatial deviations (Fig. 3f). Comparison of the five most upregulated and five most downregulated differentially expressed genes further shows that PerturbBridge generally reproduces the observed expression distributions more closely than the competing methods (Fig. 3g).

### 3.3. Ablation Study

We conduct ablation experiments on Norman to assess the contributions of the key components of PerturbBridge (Table 1). Replacing RA-AE with a conventional autoencoder consistently reduces predictive performance, supporting the benefit of explicitly preserving perturbation-associated response directions in the latent representation. Replacing PIBT with DiT leads to performance degradation across all metrics, with particularly pronounced declines in perturbation discrimination and expression prediction accuracy, highlighting the effectiveness of PIBT for conditional drift modeling. To further examine the role of condition-dependent coordinate– slot interactions, we replace the dynamic slot keys with learnable static keys while retaining condition-dependent slot representations and the read–process–write mechanism. This modification degrades performance across all metrics, suggesting that directly incorporating perturbation and bridge-time information into the coordinate–slot affinity computation provides additional benefits beyond conditioning the slot representations alone.

**Table 1.** Ablation study of PerturbBridge on the Norman dataset.

| Model | DirM ↑ | SpLFC ↑ | Pearson Δ ↑ | PDS<sub>Cos</sub> ↑ | MSE ↓ | MAE ↓ |
|---|---|---|---|---|---|---|
| Full PerturbBridge | **0.8655** | **0.8132** | **0.5621** | **0.7644** | **0.0281** | **0.0742** |
| w/o RA-AE (plain AE) | 0.8549 | 0.7931 | 0.5363 | 0.7235 | 0.0299 | 0.0746 |
| w/o PIBT (DiT) | 0.8497 | 0.7902 | 0.5284 | 0.6617 | 0.0319 | 0.0782 |
| w/ Static Slot Keys | 0.8442 | 0.7880 | 0.5213 | 0.7200 | 0.0298 | 0.0757 |

## 4. Discussion

PerturbBridge provides a conditional latent Schrödinger Bridge framework for predicting transcriptional responses to unseen perturbations from unpaired single-cell populations. By modeling perturbation-induced transitions as stochastic transport in a compact latent space, the framework avoids learning bridge dynamics directly in high-dimensional gene-expression space. RA-AE constructs a response-aware representation that explicitly preserves dominant perturbation-associated transcriptional directions while encoding complementary cellular variation. Within this latent space, PIBT parameterizes the conditional bridge drift through interactions among latent coordinates guided by perturbation conditions and bridge time.

Experiments across three genetic and chemical perturbation benchmarks demonstrate strong performance in recovering perturbation-induced transcriptional changes, particularly in differential-expression direction recovery and perturbation-effect correlation. Condition-level comparisons further indicate that these improvements extend across diverse perturbations rather than being driven by a limited number of conditions. Ablation studies further support the contributions of response-anchored representation learning and PIBT-based conditional drift modeling.

Despite these promising results, several aspects warrant further investigation. The response subspace is derived from training perturbations, and its ability to capture transcriptional response directions not adequately represented during training remains to be systematically evaluated. Moreover, the current bridge formulation adopts independent endpoint sampling as a tractable approximation, leaving the potential benefits of alternative coupling strategies unexplored. Future work will investigate more adaptive response representations and alternative coupling strategies, while extending evaluation to broader cellular contexts and pursuing prospective experimental validation.

## Data Availability

All datasets used in this study are publicly available. The Norman and Sci-Plex3 datasets were obtained from the PerturbNet data release on Hugging Face (https://huggingface.co/cyclopeta/PerturbNet_reproduce/tree/main/data_paper). The preprocessed Replogle RPE1 dataset is available through the GEARS data release on Harvard Dataverse (https://dataverse.harvard.edu/api/access/datafile/7458694).

## Funding

This work was supported by the National Natural Science Foundation of China (Grant No. 62372391), Fujian Provincial Major Science and Technology Project (Grant No. 2022YZ040011), and National Key Research and Development Program of China (Grant Nos. 2023YFF1205602, 2024YFF1206204).

## Conflict of Interest

None declared.

## Notes

### Competing Interest Statement

The authors have declared no competing interest.

### Summary of Updates

\textbf{Motivation:} Single-cell perturbation prediction seeks to recover transcriptional responses to unseen interventions from unpaired control and perturbed populations. The Schr\"odinger Bridge provides a principled framework for modeling these transitions as stochastic transport, but learning bridge dynamics directly from high-dimensional, sparse gene-expression profiles remains challenging. \textbf{Results:} We present PerturbBridge, a conditional latent Schr\"odinger Bridge framework for single-cell perturbation response prediction. A response-anchored autoencoder constructs a compact cellular representation that preserves dominant transcriptional response directions identified from training perturbations while encoding complementary cellular variation. In this latent space, PerturbBridge models conditional stochastic transport through a tractable approximation to the Schr\"odinger Bridge. We introduce the Perturbation-Induced Bridge Transformer, which uses perturbation- and time-dependent slots to guide interactions among latent coordinates during drift estimation. Evaluations on Norman, Sci-Plex3, and Replogle RPE1 demonstrate leading performance in differential-expression direction recovery and perturbation-effect correlation, alongside competitive gene-expression prediction accuracy. \textbf{Availability:} The source code is available at \url{https://github.com/echQoQ/PerturbBridge}, with an archived version at \url{https://doi.org/10.5281/zenodo.24272128}.

