## supplementary material for "PerturbBridge: Conditional Latent Schrödinger Bridge for Single-Cell Perturbation Response Prediction"

#### S1 Supplementary Implementation Details

##### S1.1 Data Preprocessing and Splitting

We use the Norman (Norman et al., 2019) and Sci-Plex3 (Srivatsan et al., 2020) datasets distributed with PerturbNet (Yu et al., 2025) and the preprocessed Replogle RPE1 dataset (Replogle et al., 2022) distributed with GEARS (Roohani et al., 2024). The supplied gene sets are retained without additional highly variable gene selection. For Norman, raw counts from the count layer are normalized to a total of $10^4$ counts per cell and transformed using $\log(1+x)$. For Sci-Plex3, the preprocessed expression matrix is used directly. For Replogle RPE1, the supplied normalized and log-transformed expression matrix is retained without further normalization. Gene ordering is preserved throughout model training, inference, and prediction export.

The resulting datasets comprise 109,738 cells and 2,279 genes for Norman, 648,737 cells and 5,087 genes for Sci-Plex3, and 162,733 cells and 5,000 genes for Replogle RPE1.

For Norman and Sci-Plex3, we follow the perturbation-level training and test splits provided with the preprocessed datasets. The Norman split holds out 95 of 236 perturbation conditions for testing. For Sci-Plex3, 38 of 188 drugs are held out, with all dose and cell-line observations associated with each test drug excluded from training. Thus, generalization is evaluated at the drug-identity level rather than at the individual drug–dose level. For Replogle RPE1, we adopt the GEARS single-perturbation splitting procedure with a random seed of 1 and nominal validation and test fractions of 0.1. The original validation partition is merged with the training partition before constructing the internal PerturbBridge validation sets, resulting in 145 held-out perturbation targets. The GEARS preprocessing procedure applies a minimum cell-count threshold of 10 cells per perturbation.

Separate internal validation sets are constructed for representation and bridge learning. For RA-AE, approximately 10% of the training perturbation identities and 10% of the control cells are reserved for validation, with control-cell sampling stratified by cell line for Sci-Plex3. For bridge learning, 20%, 10%, and 20% of the available training perturbation identities are reserved for validation on Norman, Sci-Plex3, and Replogle RPE1, respectively. Sci-Plex3 uses the same validation drug identities for both stages, whereas the genetic benchmarks use independently constructed internal validation splits. All internal splits are generated using a random seed of 42. The response basis and latent normalization statistics are estimated exclusively from the RA-AE training partition, without using expression profiles from the external test perturbations.

##### S1.2 Model Architecture and Hyperparameters

RA-AE decomposes each centered gene-expression profile into a fixed response-aligned projection and a learned complementary representation. The response basis is constructed from perturbation-induced changes in mean expression, measured relative to the corresponding control population. These response vectors are stacked as columns of a matrix $\Delta \in \mathbb{R}^{G\times M}$, with each perturbation condition contributing one vector independently of its cell count. The orthonormal basis $U \in \mathbb{R}^{G\times r}$ is obtained from the leading $r$ left singular vectors of $\Delta$. For Sci-Plex3, each response vector corresponds to a cell-line–drug–dose combination and is computed relative to the control mean of the same cell line, thereby excluding between-cell-line baseline differences from the response estimates. The centering vector $\mu$ used by RA-AE is the mean expression profile of all cells in the representation-training partition, including controls, and is distinct from the control means used to construct $\Delta$.

Let $P_\perp = I - UU^\top$ denote the projection onto the orthogonal complement of the response subspace. The unnormalized encoder and decoder are defined as

$$z_{\mathrm{raw}}(x) = \left[U^\top(x-\mu); f_\phi(P_\perp(x-\mu))\right], \tag{S1}$$

$$\widehat{x}(a,h) = \mu + Ua + P_\perp g_\psi(h), \tag{S2}$$

where $a \in \mathbb{R}^r$ contains the response-aligned coordinates and $h \in \mathbb{R}^{d-r}$ represents the complementary expression variation. The complementary encoder contains two hidden layers of widths 512 and 256, while the decoder uses hidden widths 256 and 512. All hidden layers employ GELU activations, with linear output layers and no dropout.

Following reconstruction training, the latent representation is normalized using a mean vector $m \in \mathbb{R}^d$ and a global scale factor $s$:

$$z = \frac{z_{\mathrm{raw}} - m}{s}, \qquad s = \sqrt{\frac{1}{d}\sum_{j=1}^{d}\mathrm{Var}(z_{\mathrm{raw},j})}. \tag{S3}$$

Both statistics are estimated from the representation-training partition. The decoder reverses this normalization before reconstructing gene expression. The response basis and expression centering vector remain fixed throughout RA-AE training, and the trained encoder and decoder are frozen during subsequent bridge learning.

PIBT represents each of the $d$ latent coordinates as an individual token of hidden width $H$. A set of $K$ slots is initialized from the perturbation condition and bridge time to mediate interactions among coordinate tokens. Each block constructs a shared coordinate–slot affinity matrix with projection dimension $A$. Separate softmax normalizations along the coordinate and slot dimensions enable information to be read from coordinates into slots and written back to coordinates, respectively. A slot-wise feed-forward network with an expansion ratio of 2 processes the aggregated information before the coordinate update. The condition-dependent initial slots are reused across blocks, while coordinate representations are updated sequentially. Dropout with a probability of 0.1 is applied within the PIBT blocks. Dataset-specific architecture and optimization settings are summarized in Table S1.

#### S1.3 Perturbation Condition Encoding

For genetic perturbations, we construct gene representations from the gene-to-GO annotations distributed with GEARS (Aleksander et al., 2023; Roohani et al., 2024). A binary gene–term annotation matrix is constructed after resolving gene aliases and removing annotation columns with constant values. PCA is fitted to annotated genes drawn from the union of the expression gene set and perturbation targets, using centered, unwhitened randomized PCA with seven power iterations and a random seed of 42. The resulting embedding dimensions are 256 for Norman and 384 for Replogle RPE1. Genes without available GO annotations are assigned all-zero annotation vectors before projection through the fitted PCA transformation. For combinatorial perturbations in Norman, the embeddings of the constituent target genes are summed. Although the PCA basis may include GO annotations of held-out perturbation targets, no expression measurements or perturbation-response labels from the test conditions are used to construct these embeddings.

Table S1: Architecture and optimization settings of PerturbBridge across the three benchmark datasets.

| Setting | Norman | Sci-Plex3 | Replogle RPE1 |
|---|---|---|---|
| Latent dimension $d$ | 24 | 24 | 64 |
| Response dimension $r$ | 4 | 4 | 8 |
| PIBT hidden width $H$ | 128 | 256 | 256 |
| Number of slots $K$ | 4 | 8 | 8 |
| PIBT blocks $L$ | 3 | 4 | 4 |
| Affinity dimension $A$ | 32 | 64 | 64 |
| Perturbation embedding dimension | 256 | 384 | 384 |
| RA-AE batch size | 4,096 | 4,096 | 4,096 |
| RA-AE learning rate | $5 \times 10^{-4}$ | $5 \times 10^{-4}$ | $5 \times 10^{-4}$ |
| Bridge batch size | 2,048 | 4,096 | 2,048 |
| Bridge learning rate | $10^{-4}$ | $5 \times 10^{-5}$ | $10^{-4}$ |
| State-masking annealing epochs | 50 | 50 | 10 |
| Training epochs per stage | 200 | 200 | 200 |

For chemical perturbations in Sci-Plex3, we obtain 384-dimensional drug embeddings from SMILES representations using the pretrained ChemBERTa-77M-MLM model (Chithrananda et al., 2020). The embedding of each drug is extracted from the first token of the final hidden representation, with ChemBERTa operating in evaluation mode. These pretrained embeddings remain fixed throughout bridge training. Positive drug doses are transformed using their base-10 logarithms in the units provided by the source dataset, while control doses are assigned a sentinel value of $-5$. Cell-line identities are represented as categorical features. A trainable residual multilayer perceptron adapts the fixed drug embeddings, while dose and cell-line information is incorporated through feature-wise scale and shift modulation. The dose-dependent modulation further accounts for drug identity, allowing the condition representation to vary with both treatment concentration and compound. For genetic perturbations, the residual condition encoder operates on the GO-PCA embeddings without additional dose or cell-line covariates. The downstream condition encoders are optimized jointly with PIBT.

#### S1.4 Training and Inference Settings

RA-AE is trained to minimize the mean squared reconstruction error in gene-expression space using Adam with zero weight decay. The learning rate follows a cosine schedule from an initial value of $5 \times 10^{-4}$ to zero over 200 epochs. After reconstruction training, the latent normalization statistics are estimated from the representation-training partition.

The bridge model and its condition encoder are jointly optimized using AdamW with a weight decay of $10^{-4}$. A cosine learning rate schedule is employed, with the minimum learning rate set to 1% of its initial value. Control and perturbed endpoints are sampled independently within the corresponding cellular context. For Sci-Plex3, control cells are sampled from the same cell line as the target cells. Bridge times are sampled uniformly from $\{0, 0.01, \ldots, 0.99\}$, and intermediate states are generated using a Brownian bridge with conditional variance $\sigma_{\mathrm{B}}^2 t(1-t)$, where $\sigma_{\mathrm{B}} = 1$.

To regulate the use of intermediate-state information during early training, we apply a time-dependent state-masking schedule. At epoch $e$, the intermediate latent state supplied to the drift predictor is independently replaced by a zero vector for each training example with probability

$$p_e = \frac{1}{2}\left[1 + \cos\left(\pi \min\left\{\frac{e-1}{E_{\mathrm{w}}}, 1\right\}\right)\right], \tag{S4}$$

where $E_{\mathrm{w}}$ denotes the annealing duration specified in Table S1. The bridge-matching regression target is computed from the original, unmasked latent state and remains unchanged. Perturbation conditions are not dropped during training.

We additionally employ effect-stratified sampling to adjust the representation of perturbations with different response magnitudes in training minibatches. For each perturbation condition, the effect magnitude is calculated as the Euclidean distance between its mean latent state and the corresponding control mean. For Sci-Plex3, these statistics are calculated separately for cell-line–drug–dose combinations. Perturbation conditions are assigned to four strata using the 25th, 60th, and 80th percentiles of the nonzero effect magnitudes. Target cells inherit the stratum of their corresponding perturbation condition and are sampled with replacement using relative per-cell weights of $(1, 1, 1.5, 2)$. The bridge-matching loss is then averaged over the sampled examples without additional importance weighting. For Sci-Plex3, the effect statistics exclude bridge-validation drugs. For Norman and Replogle RPE1, they are calculated from the external training partition before constructing the internal bridge-validation split.

An exponential moving average of the drift-network parameters is maintained with a decay factor of 0.99 and used for prediction. Inference is performed using Euler–Maruyama integration with 100 uniform time steps ($\Delta t = 0.01$) and a condition-guidance scale of 1. At each step, the deterministic increment is $\Delta t \, v_\theta(z_t, t, \hat{c})$. The stochastic increment has covariance $\sigma_{\mathrm{B}}^2 \Delta t I$, corresponding to a standard deviation of 0.1 per coordinate for $\sigma_{\mathrm{B}} = 1$. The final integration step is performed without stochastic noise. Negative values in the decoded expression profiles are clipped to zero before evaluation.

All PerturbBridge experiments are implemented in Python 3.7.12 and PyTorch 1.13.1 using FP32 arithmetic on a single NVIDIA GeForce RTX 3090. The released evaluation workflow uses Cell-Eval 0.7.2 (commit `44f9009`) with pdex 0.2.2 in a separate Python 3.11 environment. Each PerturbBridge configuration is trained once with a random seed of 42.

### S2 Baseline Implementation and Evaluation

#### S2.1 Baseline Implementation Details

The genetic perturbation benchmarks include CPA (Lotfollahi et al., 2023), GEARS (Roohani et al., 2024), PerturbNet (Yu et al., 2025), scGPT (Cui et al., 2024), CellFlow (Klein et al., 2025), Departures (Chi et al., 2026), and scLDM (Wu et al., 2026). For the chemical perturbation benchmark, CPA, GEARS, and scGPT are replaced by chemCPA (Hetzel et al., 2022) and PRnet (Qi et al., 2024). A Control baseline is included to assess prediction performance in the absence of a learned perturbation response.

Each baseline follows its method-specific representation and optimization settings. For genetic perturbations, CPA uses GO-PCA perturbation embeddings and a Gaussian expression decoder, with predictions obtained from the conditional decoder means without additional sampling from the predicted variance. CellFlow uses 50-dimensional PCA representations fitted to its training cells and controls. scLDM employs the released gene2vec embeddings for genetic perturbations and ChemBERTa features with a scaled log-dose covariate for Sci-Plex3. Its cellular representations are obtained from a frozen variational autoencoder, with control–target matching performed within the corresponding perturbation condition and cellular context.

The comparison combines dataset-specific baseline reproductions with existing prediction exports for Norman (Departures, GEARS, PerturbNet, and scGPT) and Sci-Plex3 (chemCPA, PRnet, and PerturbNet). Training budgets and internal validation procedures follow the respective baseline configurations. All numerical comparisons are performed in the retained gene-expression space using consistent Cell-Eval metric definitions.

#### S2.2 Evaluation Metrics

We evaluate perturbation responses using eight metrics from Cell-Eval version 0.7.2 (Adduri et al., 2026). Let $x_{pig}$ and $\widehat{x}_{pjg}$ denote the observed and predicted log-normalized expression of gene $g$ in cells $i$ and $j$ under perturbation $p$, respectively. With $n_p$ observed cells and $\widehat{n}_p$ predicted cells, the pseudobulk profiles are

$$\mu_{pg} = \frac{1}{n_p}\sum_{i=1}^{n_p} x_{pig}, \qquad \widehat{\mu}_{pg} = \frac{1}{\widehat{n}_p}\sum_{j=1}^{\widehat{n}_p} \widehat{x}_{pjg}. \tag{S5}$$

Let $\mu_0$ and $\widehat{\mu}_0$ denote the corresponding control profiles. The observed and predicted perturbation effects are

$$\Delta_p = \mu_p - \mu_0, \qquad \widehat{\Delta}_p = \widehat{\mu}_p - \widehat{\mu}_0. \tag{S6}$$

For PerturbBridge, the same observed control cells are included in both the prediction and reference exports. For Sci-Plex3, the overall benchmark groups cells by drug identity, pooling across doses and cell lines; dose-stratified analyses are evaluated separately.

**Expression accuracy.** Mean squared error (MSE) and mean absolute error (MAE) measure pseudobulk prediction error across the $G$ evaluated genes:

$$\mathrm{MSE}_p = \frac{1}{G}\sum_{g=1}^{G}(\widehat{\mu}_{pg} - \mu_{pg})^2, \tag{S7}$$

$$\mathrm{MAE}_p = \frac{1}{G}\sum_{g=1}^{G}|\widehat{\mu}_{pg} - \mu_{pg}|. \tag{S8}$$

Pearson $\Delta$ measures the correlation between the predicted and observed expression changes:

$$\text{Pearson}\,\Delta_p = \frac{\sum_{g=1}^{G}(\Delta_{pg} - \overline{\Delta}_p)(\widehat{\Delta}_{pg} - \overline{\widehat{\Delta}}_p)}{\sqrt{\sum_{g=1}^{G}(\Delta_{pg} - \overline{\Delta}_p)^2}\sqrt{\sum_{g=1}^{G}(\widehat{\Delta}_{pg} - \overline{\widehat{\Delta}}_p)^2}}, \tag{S9}$$

where $\overline{\Delta}_p = G^{-1}\sum_g \Delta_{pg}$ and $\overline{\widehat{\Delta}}_p = G^{-1}\sum_g \widehat{\Delta}_{pg}$.

**Differential-expression recovery.** Differential expression is computed against the corresponding control cells using the Mann–Whitney $U$ test implemented in pdex, with Benjamini–Hochberg correction across genes for each comparison. Let $q_{pg}$ denote the adjusted observed-data $p$-value and define

$$\mathcal{S}_p = \{g : q_{pg} < 0.05\}. \tag{S10}$$

With log1p-transformed input, the default geometric-mean convention in pdex gives the following back-transformed summaries and $\log_2$ fold changes:

$$b_{pg} = \exp(\mu_{pg}) - 1, \qquad \widehat{b}_{pg} = \exp(\widehat{\mu}_{pg}) - 1, \tag{S11}$$

$$\ell_{pg} = \log_2 \frac{b_{pg}}{b_{0g}}, \qquad \widehat{\ell}_{pg} = \log_2 \frac{\widehat{b}_{pg}}{\widehat{b}_{0g}}. \tag{S12}$$

Here $b_{0g}$ and $\widehat{b}_{0g}$ are defined analogously for controls; the default pseudocount is zero. DE Direction Match (DirM) measures sign agreement on genes significant in the observed data:

$$\mathrm{DirM}_p = \frac{1}{|\mathcal{S}_p|}\sum_{g\in\mathcal{S}_p} \mathbf{1}\left[\mathrm{sgn}(\ell_{pg}) = \mathrm{sgn}(\widehat{\ell}_{pg})\right]. \tag{S13}$$

Spearman correlation of log-fold changes (SpLFC) measures agreement between their rankings over the same gene set. Let $R_{pg}$ and $\widehat{R}_{pg}$ be the ranks of $\ell_{pg}$ and $\widehat{\ell}_{pg}$ within $\mathcal{S}_p$, assigning average ranks to ties, and let $\overline{R}_p$ and $\overline{\widehat{R}}_p$ denote their respective means. Then

$$\mathrm{SpLFC}_p = \frac{\sum_{g\in\mathcal{S}_p}(R_{pg}-\overline{R}_p)(\widehat{R}_{pg}-\overline{\widehat{R}}_p)}{\sqrt{\sum_{g\in\mathcal{S}_p}(R_{pg}-\overline{R}_p)^2}\sqrt{\sum_{g\in\mathcal{S}_p}(\widehat{R}_{pg}-\overline{\widehat{R}}_p)^2}}. \tag{S14}$$

These correspond to the `de_direction_match` and `de_spearman_lfc_sig` outputs of Cell-Eval, respectively.

**Perturbation discrimination.** Perturbation Discrimination Score (PDS) evaluates whether each predicted effect is closest to its corresponding observed effect among the $M$ evaluated perturbations. Following Cell-Eval, let $\mathcal{G}_p$ contain the evaluated genes except those whose names exactly match the perturbation label $p$. Composite labels therefore do not individually exclude their constituent target genes. For $a = \widehat{\Delta}_p|_{\mathcal{G}_p}$ and $b = \Delta_q|_{\mathcal{G}_p}$, the three distance functions are

$$d_{L_1}(a,b) = \sum_{g\in\mathcal{G}_p} |a_g - b_g|, \tag{S15}$$

$$d_{L_2}(a,b) = \sqrt{\sum_{g\in\mathcal{G}_p} (a_g - b_g)^2}, \tag{S16}$$

$$d_{\cos}(a,b) = 1 - \frac{\sum_{g\in\mathcal{G}_p} a_g b_g}{\sqrt{\sum_{g\in\mathcal{G}_p} a_g^2}\sqrt{\sum_{g\in\mathcal{G}_p} b_g^2}}. \tag{S17}$$

When either input vector has zero norm, the cross-distance implementation assigns cosine distance 1. For each distance $d$, let $\pi_p^{(d)}$ be the permutation of candidate perturbations obtained by sorting $d(\widehat{\Delta}_p, \Delta_q)$ in ascending order, with both vectors restricted to $\mathcal{G}_p$. The zero-based rank of the matching perturbation and its score are

$$r_p^{(d)} = \left(\pi_p^{(d)}\right)^{-1}(p) - 1, \qquad \mathrm{PDS}_p^{(d)} = 1 - \frac{r_p^{(d)}}{M}. \tag{S18}$$

The permutation positions are numbered from 1 to $M$; ties follow the ordering returned by NumPy's `argsort`, as in Cell-Eval. We report $\mathrm{PDS}_{L_1}$, $\mathrm{PDS}_{L_2}$, and $\mathrm{PDS}_{\cos}$, each ranging from $1/M$ to 1.

**Aggregation.** For metric $m$, let $\mathcal{P}_m$ be the set of perturbations with valid scores. The reported mean is

$$\overline{m} = \frac{1}{|\mathcal{P}_m|}\sum_{p\in\mathcal{P}_m} m_p. \tag{S19}$$

Undefined correlations are excluded from aggregation. Lower MSE and MAE indicate better expression accuracy; higher DirM, SpLFC, Pearson $\Delta$, and PDS indicate better predictive performance. Box plots display the distributions of per-perturbation scores across the evaluated test conditions.

### S3 Additional Experimental Details

#### S3.1 Complete Benchmark Results

Complete benchmark results for Norman, Sci-Plex3, and Replogle RPE1 are presented in Tables S2–S4. The best result for each metric is highlighted in bold.

Table S2: Complete benchmark results on Norman.

| Method | DirM ↑ | SpLFC ↑ | Pearson Δ ↑ | $PDS_{L_1}$ ↑ | $PDS_{L_2}$ ↑ | $PDS_{Cos}$ ↑ | MSE ↓ | MAE ↓ |
|---|---|---|---|---|---|---|---|---|
| Control | 0.4360 | -0.2317 | 0.0177 | 0.5092 | 0.5079 | 0.5525 | 0.0465 | 0.0898 |
| CPA | 0.5795 | 0.3154 | 0.2959 | 0.5022 | 0.5022 | 0.5084 | 0.0812 | 0.1233 |
| GEARS | 0.7885 | 0.7597 | 0.3160 | 0.5783 | 0.5797 | 0.6153 | 0.0584 | 0.1646 |
| PerturbNet | 0.6131 | 0.0912 | 0.3616 | 0.5087 | 0.5033 | 0.4940 | 0.0391 | 0.0837 |
| scGPT | 0.8298 | 0.6867 | 0.4984 | 0.6052 | 0.6515 | 0.7432 | 0.0396 | 0.0944 |
| CellFlow | 0.8022 | 0.6543 | 0.4728 | 0.6716 | 0.6778 | 0.7091 | 0.0311 | **0.0733** |
| Departures | 0.6346 | 0.2129 | 0.3629 | 0.5206 | 0.5345 | 0.5747 | 0.0455 | 0.1029 |
| scLDM | 0.4710 | -0.3464 | 0.3156 | 0.5736 | 0.5827 | 0.6588 | 0.0468 | 0.0989 |
| PerturbBridge | **0.8655** | **0.8132** | **0.5621** | **0.6983** | **0.7179** | **0.7644** | **0.0281** | 0.0742 |

Table S3: Complete benchmark results on Sci-Plex3.

| Method | DirM ↑ | SpLFC ↑ | Pearson Δ ↑ | $PDS_{L_1}$ ↑ | $PDS_{L_2}$ ↑ | $PDS_{Cos}$ ↑ | MSE ↓ | MAE ↓ |
|---|---|---|---|---|---|---|---|---|
| Control | 0.5023 | 0.0047 | 0.0434 | 0.5132 | 0.5132 | 0.5132 | 0.0037 | 0.0274 |
| chemCPA | 0.7724 | 0.3849 | 0.5967 | **0.7479** | **0.7701** | 0.8068 | 0.0036 | 0.0314 |
| PRnet | 0.6122 | 0.4168 | 0.4101 | 0.6219 | 0.6517 | 0.6898 | 0.0225 | 0.1309 |
| PerturbNet | 0.7975 | 0.6158 | 0.6406 | 0.7209 | 0.7368 | 0.8012 | 0.0025 | 0.0234 |
| CellFlow | 0.8256 | 0.6253 | 0.5687 | 0.6974 | 0.7611 | 0.7756 | 0.0035 | 0.0270 |
| Departures | 0.7294 | 0.6842 | 0.5518 | 0.6904 | 0.7521 | 0.7632 | 0.0051 | 0.0442 |
| scLDM | 0.5684 | 0.0321 | 0.2955 | 0.7036 | 0.7285 | 0.7936 | 0.0096 | 0.0590 |
| PerturbBridge | **0.8705** | **0.7099** | **0.6641** | 0.7431 | 0.7652 | **0.8172** | **0.0024** | **0.0215** |

Table S4: Complete benchmark results on Replogle RPE1.

| Method | DirM ↑ | SpLFC ↑ | Pearson Δ ↑ | $PDS_{L_1}$ ↑ | $PDS_{L_2}$ ↑ | $PDS_{Cos}$ ↑ | MSE ↓ | MAE ↓ |
|---|---|---|---|---|---|---|---|---|
| Control | 0.4925 | -0.0211 | -0.0041 | 0.5034 | 0.5037 | 0.4848 | 0.0216 | 0.0898 |
| CPA | 0.6104 | 0.2528 | 0.1857 | 0.5049 | 0.5063 | 0.5346 | 0.0181 | 0.0790 |
| GEARS | 0.8199 | 0.6391 | 0.5574 | 0.5810 | 0.5910 | 0.5541 | 0.0115 | 0.0687 |
| PerturbNet | 0.7490 | 0.4958 | 0.4595 | 0.5044 | 0.5044 | 0.4953 | 0.0153 | 0.0797 |
| scGPT | 0.8278 | **0.6572** | 0.5823 | 0.5042 | 0.5035 | 0.5033 | 0.0123 | 0.0691 |
| CellFlow | 0.8358 | 0.6406 | 0.5892 | 0.6378 | 0.6443 | 0.5922 | 0.0112 | 0.0649 |
| Departures | 0.7314 | 0.4334 | 0.4251 | 0.5094 | 0.5135 | 0.5018 | 0.0166 | 0.0852 |
| scLDM | 0.7586 | 0.4244 | 0.5077 | 0.5375 | 0.5405 | 0.5175 | 0.0174 | 0.0905 |
| PerturbBridge | **0.8403** | 0.6481 | **0.5967** | **0.6717** | **0.6834** | **0.6055** | **0.0106** | **0.0644** |

##### S3.2 Ablation Study Design

We conduct three ablation experiments on Norman to investigate the contributions of response-anchored representation learning, the conditional drift architecture, and condition-dependent coordinate–slot affinities. All variants follow the same data-splitting protocol, bridge-matching objective, effect-stratified sampling procedure, and state-masking schedule as the full model. The GO-PCA perturbation embeddings are also kept unchanged. All bridge variants are trained for 200 epochs with a random seed of 42. Predictions use exponential moving average parameters and 100 stochastic integration steps. Evaluation follows Section S2, and the results are presented in the ablation table of the main text.

**Replacing RA-AE with a conventional autoencoder.** The *w/o RA-AE (plain AE)* variant replaces the response-anchored representation with a conventional deterministic multilayer perceptron autoencoder. The encoder has dimensions $G \to 512 \to 256 \to 24$, while the decoder follows $24 \to 256 \to 512 \to G$. All hidden layers use GELU activations, with linear output layers. Unlike RA-AE, the plain autoencoder learns all 24 latent coordinates solely through reconstruction, without a fixed response projection or an orthogonal residual decomposition. It follows the same representation-training protocol as RA-AE, including data partitions, reconstruction objective, and optimization settings. Latent normalization statistics are estimated from its representation-training partition using the same procedure as the full model. After training, the autoencoder is frozen, and a new PIBT model is trained using the original bridge architecture and optimization settings. This comparison isolates the contribution of response-anchored representation learning while maintaining the same latent dimension.

**Replacing PIBT with a DiT backbone.** The *w/o PIBT (DiT)* variant retains the pretrained RA-AE representation but replaces PIBT with a conditional Diffusion Transformer (DiT) architecture (Peebles and Xie, 2023). Each of the 24 latent coordinates is represented as an individual token. The backbone comprises three Transformer blocks with a hidden width of 96, three attention heads, a feed-forward expansion ratio of 2, and a dropout probability of 0.1. Full multi-head self-attention enables direct interactions among coordinate tokens, followed by token-wise feed-forward transformations. Perturbation and time embeddings condition the Transformer blocks through adaptive layer normalization with zero-initialized residual gates (adaLN-Zero). A shared scalar prediction head, initialized to zero, maps each coordinate representation to its corresponding drift component. The perturbation-condition encoder and bridge-training procedure remain unchanged. This ablation compares PIBT with a conventional conditional Transformer alternative for drift parameterization.

**Replacing condition-dependent slot keys with static keys.** The *w/ Static Slot Keys* variant retains the pretrained RA-AE representation and the original PIBT configuration $(H, K, L, A) = (128, 4, 3, 32)$. The only modification concerns the construction of the affinity keys. Let $I_{\text{slot}}$ denote the learnable slot identity embeddings. In block $\ell$, the slot keys are computed as

$$K_s^\ell = \text{LN}(I_{\text{slot}})W_K^\ell, \tag{S20}$$

rather than being projected from the perturbation- and time-conditioned slots $S^{(0)}$. The coordinate queries, shared affinity matrix, and both softmax normalizations remain unchanged. Importantly, the conditioned slots $S^{(0)}$ are still used in the read–process–write pathway, allowing perturbation and time information to influence the coordinate updates. Consequently, the affinity keys are independent of the perturbation condition and bridge time, while the affinities remain dependent on the evolving coordinate representations. This comparison evaluates the contribution of directly conditioning the affinity keys, beyond incorporating conditional information into the slot representations.

### S4 Additional Theoretical Details

#### S4.1 Conditional Reference Bridge Construction

We consider a fixed cellular context $\tau$ and perturbation condition $c$, with latent control and perturbed distributions $\rho_0^\tau$ and $\rho_1^{\tau,c}$, respectively. As described in the main text, PerturbBridge adopts a Brownian reference process

$$dZ_t = \sigma_\mathrm{B}\, dW_t, \qquad Z_0 \sim \rho_0^\tau, \tag{S21}$$

where $\sigma_\mathrm{B} > 0$ is the constant diffusion scale. The transition density over an interval of length $s$ is

$$k_s(z' \mid z) = \frac{1}{(2\pi\sigma_\mathrm{B}^2 s)^{d/2}} \exp\left(-\frac{\|z' - z\|_2^2}{2\sigma_\mathrm{B}^2 s}\right). \tag{S22}$$

Conditioning the reference process on $Z_0 = z_0$ and $Z_1 = z_1$ yields a Brownian bridge. Its intermediate distribution can be expressed as

$$q(z_t \mid z_0, z_1) = \mathcal{N}\left(z_t;\, (1-t)z_0 + tz_1,\, \sigma_\mathrm{B}^2 t(1-t) I_d\right), \tag{S23}$$

for $0 < t < 1$. This conditional Gaussian distribution provides the analytic sampling rule used in PerturbBridge, following the reference bridge construction employed in I$^2$SB (Liu et al., 2023).

The corresponding endpoint-conditioned drift is obtained from the transition density of the reference process:

$$\begin{aligned} b_\mathrm{ref}(z, z_1, t) &= \sigma_\mathrm{B}^2 \nabla_z \log k_{1-t}(z_1 \mid z) \\ &= \frac{z_1 - z}{1-t}. \end{aligned} \tag{S24}$$

Consequently, the reference process conditioned on the terminal state satisfies

$$dZ_t = b_\mathrm{ref}(Z_t, z_1, t)\, dt + \sigma_\mathrm{B}\, dW_t. \tag{S25}$$

In the Schrödinger Bridge formulation, the optimal path measure is obtained by combining conditional reference bridges under an entropy-minimizing endpoint coupling (Léonard, 2013). PerturbBridge instead uses independently sampled endpoints, corresponding to

$$\gamma^{\tau,c} = \rho_0^\tau \otimes \rho_1^{\tau,c}. \tag{S26}$$

Let $\mathbb{R}^\tau$ denote the reference path measure. The resulting mixture of conditional reference bridges is

$$\mathbb{Q}^{\tau,c} = \int \mathbb{R}^\tau(\cdot \mid z_0, z_1)\, \gamma^{\tau,c}(dz_0, dz_1). \tag{S27}$$

Because the endpoint coupling has marginals $\rho_0^\tau$ and $\rho_1^{\tau,c}$, the bridge mixture satisfies the prescribed endpoint constraints. The independent coupling is not generally entropy-optimal, and $\mathbb{Q}^{\tau,c}$ therefore need not coincide with the optimal Schrödinger Bridge path measure.

#### S4.2 Marginal Preservation under Bridge Matching

PerturbBridge learns a Markovian drift field from the conditional reference bridges through the bridge-matching objective (Shi et al., 2023). For a drift function $v$, the unmasked population objective is

$$\mathcal{L}_\mathrm{SB}(v) = \mathbb{E}\left[\frac{1}{d}\left\|v(Z_t, t, \tau, c) - b_\mathrm{ref}(Z_t, Z_1, t)\right\|_2^2\right], \tag{S28}$$

where the expectation is taken over cellular contexts, perturbation conditions, bridge times, independently sampled endpoint pairs, and intermediate states drawn from the corresponding conditional reference bridges. The analysis considers the continuous-time extension of the regression problem after state masking has been annealed to zero. For the squared-loss statements, time sampling is restricted to $[0, T]$ with $T < 1$ and finite second moments are assumed; this avoids the singular terminal drift. The implementation uses the discrete time grid specified in Section S1.

By the conditional expectation property of squared-error regression, the population minimizer is

$$v^*(z, t, \tau, c) = \mathbb{E}_{\mathbb{Q}^{\tau,c}}\left[b_{\mathrm{ref}}(Z_t, Z_1, t) \mid Z_t = z\right]. \tag{S29}$$

More precisely, for square-integrable drift functions, the objective admits the decomposition

$$\begin{aligned}\mathcal{L}_{\mathrm{SB}}(v) = \mathcal{L}_{\mathrm{SB}}(v^*)& \\ + \frac{1}{d}\mathbb{E}&\left[\|v(Z_t, t, \tau, c) - v^*(Z_t, t, \tau, c)\|_2^2\right].\end{aligned} \tag{S30}$$

Thus, matching the endpoint-conditioned reference drift recovers its conditional expectation in the population limit.

To examine the resulting marginal dynamics, let $p_t^{\tau,c}$ denote the time-$t$ marginal density of $\mathbb{Q}^{\tau,c}$. For a smooth, compactly supported test function $\varphi$, applying Itô's formula to the conditional reference bridges and averaging over their endpoint distribution gives

$$\begin{aligned}\frac{d}{dt}\mathbb{E}_{\mathbb{Q}^{\tau,c}}[\varphi(Z_t)] &= \mathbb{E}_{\mathbb{Q}^{\tau,c}}\left[b_{\mathrm{ref}}(Z_t, Z_1, t) \cdot \nabla\varphi(Z_t)\right] \\ &\quad + \frac{\sigma_{\mathrm{B}}^2}{2}\mathbb{E}_{\mathbb{Q}^{\tau,c}}[\Delta\varphi(Z_t)].\end{aligned} \tag{S31}$$

Using the definition of $v^*$ and the law of total expectation, we obtain

$$\begin{aligned}\frac{d}{dt}\int \varphi(z) p_t^{\tau,c}(z)\, dz &= \int v^*(z, t, \tau, c) \cdot \nabla\varphi(z)\, p_t^{\tau,c}(z)\, dz \\ &\quad + \frac{\sigma_{\mathrm{B}}^2}{2}\int \Delta\varphi(z)\, p_t^{\tau,c}(z)\, dz.\end{aligned} \tag{S32}$$

Equivalently, the marginal density satisfies the Fokker–Planck equation in the weak sense:

$$\frac{\partial p_t^{\tau,c}}{\partial t} = -\nabla \cdot \left(p_t^{\tau,c} v^*(\cdot, t, \tau, c)\right) + \frac{\sigma_{\mathrm{B}}^2}{2}\Delta p_t^{\tau,c}. \tag{S33}$$

Consider the Markovian diffusion

$$d\widetilde{Z}_t = v^*(\widetilde{Z}_t, t, \tau, c)\, dt + \sigma_{\mathrm{B}}\, dW_t, \qquad \widetilde{Z}_0 \sim \rho_0^{\tau}. \tag{S34}$$

Its marginal distributions satisfy the same Fokker–Planck equation and initial condition as those of the reference bridge mixture. Under appropriate regularity and uniqueness conditions, the two processes therefore share the same time marginals, consistent with Markovian projection (Gyöngy, 1986; Shi et al., 2023). With the corresponding endpoint limits, the terminal distribution of the Markovian diffusion is $\rho_1^{\tau,c}$.

This correspondence concerns time marginals rather than the full path measure or endpoint coupling. In particular, it does not imply entropy optimality under the independently sampled endpoint coupling. PerturbBridge approximates the population drift using PIBT and performs inference through finite-step stochastic integration. Consequently, exact marginal preservation is a property of the idealized continuous-time formulation rather than a guarantee for the learned finite-sample model.

#### Declaration of AI-Assisted Tools

We used OpenAI ChatGPT (GPT-6) and DeepSeek (DeepSeek-V4.1-Flash) to assist with code development and debugging. We independently reviewed and validated the AI-assisted code. All research ideas, methodological contributions, experimental analyses, and scientific conclusions presented in this study are our own work, and we take full responsibility for the accuracy and integrity of the results.

Seyone Chithrananda, Gabriel Grand, and Bharath Ramsundar. Chemberta: large-scale self-supervised pretraining for molecular property prediction. *arXiv preprint arXiv:2010.09885*, 2020.

Haotian Cui, Chloe Wang, Hassaan Maan, Kuan Pang, Fengning Luo, Nan Duan, and Bo Wang. scgpt: toward building a foundation model for single-cell multi-omics using generative ai. *Nature methods*, 21(8):1470–1480, 2024.

István Gyöngy. Mimicking the one-dimensional marginal distributions of processes having an itô differential. *Probability theory and related fields*, 71(4):501–516, 1986.

Leon Hetzel, Simon Boehm, Niki Kilbertus, Stephan Günnemann, Fabian Theis, et al. Predicting cellular responses to novel drug perturbations at a single-cell resolution. *Advances in Neural Information Processing Systems*, 35:26711–26722, 2022.

Dominik Klein, Jonas Simon Fleck, Daniil Bobrovskiy, Lea Zimmermann, Sören Becker, Alessandro Palma, Leander Dony, Alejandro Tejada-Lapuerta, Guillaume Huguet, Hsiu-Chuan Lin, et al. Cellflow enables generative single-cell phenotype modeling with flow matching. *bioRxiv*, pages 2025–04, 2025.

Christian Léonard. A survey of the schrödinger problem and some of its connections with optimal transport. *arXiv preprint arXiv:1308.0215*, 2013.

Guan-Horng Liu, Arash Vahdat, De-An Huang, Evangelos A Theodorou, Weili Nie, and Anima Anandkumar. I2SB: Image-to-image Schrödinger bridge. *arXiv preprint arXiv:2302.05872*, 2023.

Mohammad Lotfollahi, Anna Klimovskaia Susmelj, Carlo De Donno, Leon Hetzel, Yuge Ji, Ignacio L Ibarra, Sanjay R Srivatsan, Mohsen Naghipourfar, Riza M Daza, Beth Martin, et al. Predicting cellular responses to complex perturbations in high-throughput screens. *Molecular systems biology*, 19(6):MSB202211517, 2023.

Thomas M Norman, Max A Horlbeck, Joseph M Replogle, Alex Y Ge, Albert Xu, Marco Jost, Luke A Gilbert, and Jonathan S Weissman. Exploring genetic interaction manifolds constructed from rich single-cell phenotypes. *Science*, 365(6455):786–793, 2019.

William Peebles and Saining Xie. Scalable diffusion models with transformers. In *2023 IEEE/CVF International Conference on Computer Vision (ICCV)*, pages 4172–4182. IEEE, 2023.

Xiaoning Qi, Lianhe Zhao, Chenyu Tian, Yueyue Li, Zhen-Lin Chen, Peipei Huo, Runsheng Chen, Xiaodong Liu, Baoping Wan, Shengyong Yang, et al. Predicting transcriptional responses to novel chemical perturbations using deep generative model for drug discovery. *Nature Communications*, 15(1):9256, 2024.

Joseph M Replogle, Reuben A Saunders, Angela N Pogson, Jeffrey A Hussmann, Alexander Lenail, Alina Guna, Lauren Mascibroda, Eric J Wagner, Karen Adelman, Gila Lithwick-Yanai, et al. Mapping information-rich genotype-phenotype landscapes with genome-scale perturb-seq. *Cell*, 185(14):2559–2575, 2022.

Yusuf Roohani, Kexin Huang, and Jure Leskovec. Predicting transcriptional outcomes of novel multigene perturbations with gears. *Nature biotechnology*, 42(6):927–935, 2024.

Yuyang Shi, Valentin De Bortoli, Andrew Campbell, and Arnaud Doucet. Diffusion schrödinger bridge matching. *Advances in neural information processing systems*, 36:62183–62223, 2023.

Sanjay R Srivatsan, José L McFaline-Figueroa, Vijay Ramani, Lauren Saunders, Junyue Cao, Jonathan Packer, Hannah A Pliner, Dana L Jackson, Riza M Daza, Lena Christiansen, et al. Massively multiplex chemical transcriptomics at single-cell resolution. *Science*, 367(6473): 45–51, 2020.

Boyang Wu, Yuhang Liu, Yue Cheng, Xiangrong Liu, Quan Zou, Leyi Wei, and Lei Xu. scldm: a conditional diffusion framework for single-cell perturbation prediction. *Bioinformatics*, 42 (10):btag727, 2026.

Hengshi Yu, Weizhou Qian, Yuxuan Song, and Joshua D Welch. Perturbnet predicts single-cell responses to unseen chemical and genetic perturbations. *Molecular systems biology*, 21(8): 960–982, 2025.
